# Estimating orientation in Natural scenes: A Spiking Neural Network Model of the Insect Central Complex

**DOI:** 10.1101/2024.02.13.580047

**Authors:** Rachael Stentiford, James Knight, Thomas Nowotny, Andrew Philippides, Paul Graham

## Abstract

The central complex of insects contains cells, organised as a ring attractor, that encode head direction. The ‘bump’ of activity in the ring can be updated by idiothetic cues and external sensory information. Plasticity at the synapses between these cells and the ring neurons, that are responsible for bringing sensory information into the central complex, has been proposed to form a mapping between visual cues and the heading estimate which allows for more accurate tracking of the current heading, than if only idiothetic information were used.

In Drosophila, ring neurons have well characterised non-linear receptive fields. In this work we produce synthetic versions of these visual receptive fields using a combination of excitatory inputs and mutual inhibition between ring neurons. We use these receptive fields to bring visual information into a spiking neural network model of the insect central complex based on the recently published Drosophila connectome.

Previous modelling work has focused on how this circuit functions as a ring attractor using the same type of simple visual cues commonly used experimentally. While we initially test the model on these simple stimuli, we then go on to apply the model to complex natural scenes containing multiple conflicting cues. We show that this simple visual filtering provided by the ring neurons is sufficient to form a mapping between heading and visual features and maintain the heading estimate in the absence of angular velocity input. The network is successful at tracking heading even when presented with videos of natural scenes containing conflicting information from environmental changes and translation of the camera.

**Author summary:** To navigate through the world animals require knowledge of the direction they are facing. Insects keep track of this “head direction” with a population of ‘compass’ neurons. These cells can use internal measures of angular velocity to maintain the heading estimate but this becomes inaccurate over time and needs to be stabilised by environmental cues. In this work we produce a spiking neural network model replicating the connectivity between regions of the insect brain known to be involved in keeping track of heading and the neurons which are responsible for bringing sensory information into the circuit.

We show that the model replicates the dynamics of visual learning from experiments where flies learn simple visual stimuli. Then using panoramic videos of complex natural environments, we show that the learned mapping between the current estimate of heading in the compass neurons and the features of the visual scene can maintain and enforce the correct heading estimate.

## Introduction

Maintaining a stable estimate of heading direction is essential for many behaviours across species, including complex navigation behaviours such as path integration which enables animals to return directly to the nest after taking an indirect outward path [1–3]. In insects, heading direction is tracked by ‘compass neurons’ (EPG cells) in the ellipsoid body (EB) of the central complex that show activity with strong directional tuning which can be tied to visual features in the environment [4] analogous to head direction cells reported in mammals [5, 6]. Unlike in mammals where head direction cells are distributed across several brain regions, dendrites of insect EPG neurons are arranged in a ring (*Drosophila*, [4]) or arc (Locust, [7]; Bee, [8]) within the EB, with adjacent neurons tuned to adjacent heading directions. There have been several previous computational models describing this circuit functioning as a ring attractor, but none of them have been tested using natural stimuli or include bioplausible filtering of the visual input. There has also been a lot of experimental work on ring neurons, in particular the visual receptive fields of two subypes ER2 and ER4d (referred to as R2 and R4 populations in our model) which are known to deliver sensory input to the EB. This work brings together visual processing by ring neurons with *Drosophila* like receptive fields, and a spiking neural network model of the central complex ring attractor network challenged with natural visual stimuli.

Fig 1B shows the model structure comprised of 6 populations of cells, excitatory EPG and PEN cells, and inhibitory R, Δ7, R2 and R4 ring neurons. Visual input to the model arrives via the R2 and R4 ring neuron populations and PEN neurons encode angular velocity (the ideothetic estimate of heading change). The mapping between visual information and the heading representation is learnt at synapses between Ring and EPG neurons. We use the activity of the EPG cells as the model output and test the accuracy of the model by observing how well these cells estimate the ground truth heading. We test the model first on visual stimuli frequently used in experimental studies before challenging the network with natural visual scenes containing conflicting information.

**Fig 1.**
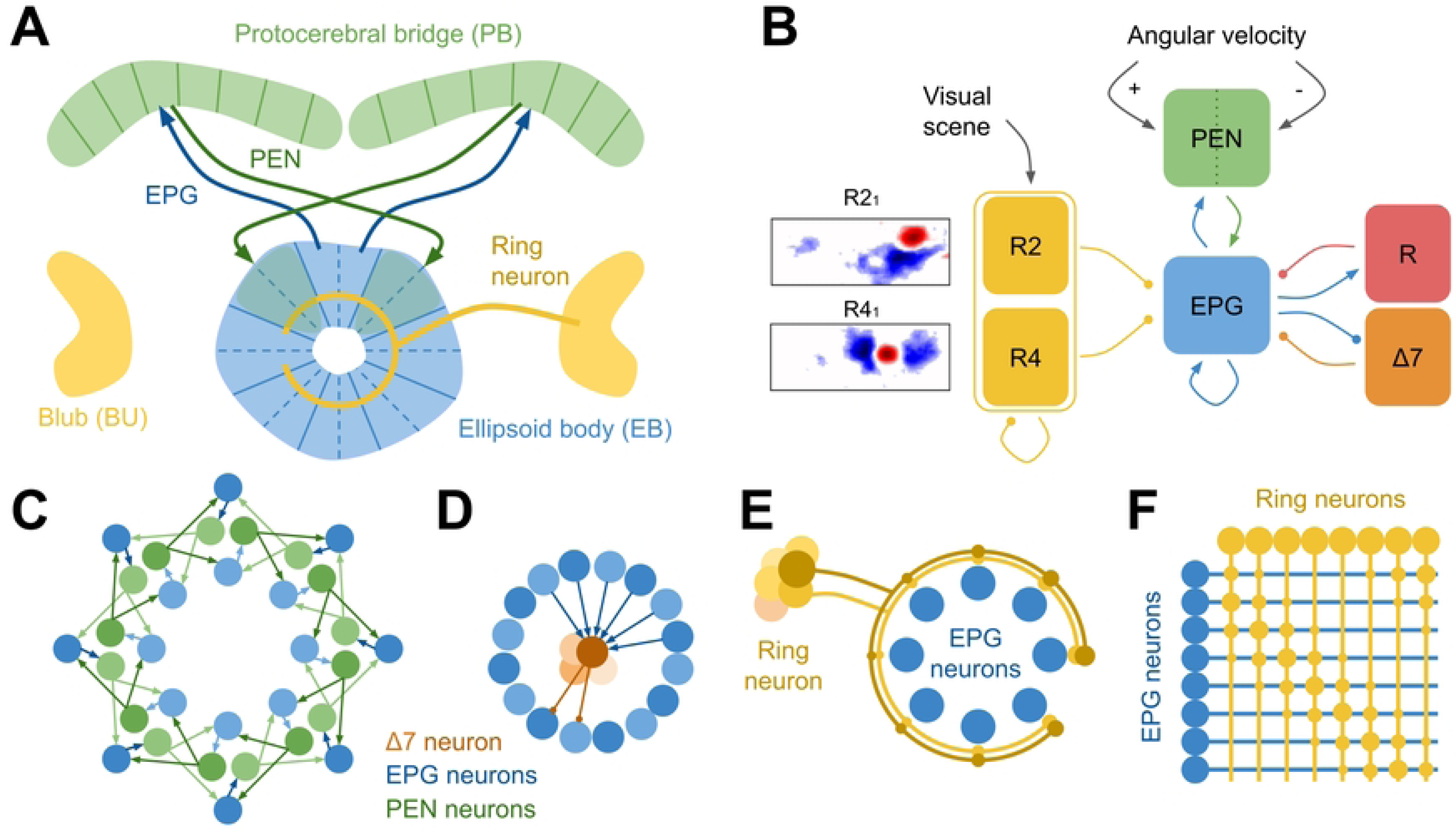
Organisation of head direction circuit in the Central Complex. (A) Major central complex regions responsible for heading tracking: ring shaped ellipsoid body (EB; blue), bulb (BU; yellow) and protocerebral bridge (PB; green). Projection patterns of two example EPG cells (blue arrows) each from one EB wedge projecting to the equivalent glomeruli (left or right) of the PB; and PEN cells (green arrows) returning projections to one tile (made up of 2 wedges) clockwise or anticlockwise further around the ring. B: Six populations of Leaky Integrate and Fire (LIF) neurons representing cell types of the central complex. Angular velocity information is provided to PEN neurons (green), positive and negative angular velocities are delivered to the right and left PEN neurons respectively. Visual scenes are provided via R2 and R4 ring neurons (yellow). Ring neuron activation is determined by each cell’s receptive field (two examples of averaged Drosophila ring neuron receptive fields [9] are shown, with excitatory (red) and inhibitory (blue) regions and the field of view indicated by the frame, which is 120 x 270 degrees). EPG cells (blue) send excitatory connections back onto themselves and to R (red), Δ7 (orange), and PEN cells. EPG cells receive excitatory input from PEN cells, and inhibitory connections from R, Δ7, R2 and R4 cells. C: Effective circuit connections between each of 16 EPG (blue) and 16 PEN (green) cells split into right and left projecting cells. PEN cells return projections to both EPG cells one wedge further clockwise or anticlockwise around the ringD: Connections between all EPG cells and a single (out of 8) Δ7 cell. The connection pattern results in strongest inhibitory input to EPG cells opposite to the bump position. E: Each ring neuron makes connections with every EPG cell in the ring. F: Matrix showing all ring neuron to EPG cell connections, including strengthened weights between coactive ring and EPG cells.

Due the shape of the EB and the topographical arrangement of head direction cells, calcium imaging of the EB can reveal a single ‘bump’ of activity advances around the ring structure whilst the heading of the insect changes. Additional to the EB, the insect central complex contains several neuropils including the protocerebral bridge (PB; Fig 1A). PEN neurons (specifically PEN-a neurons) in the PB also show directional turning, and conjunctively encode heading and angular velocity, producing two sustained bumps of activity within the PB, one per hemisphere [10]; [11]. These activity patterns and the known connectivity within the central complex [12–14] strongly support ring attractor dynamics similar to those in the mammalian head direction system [15–19]. Ring attractors function as winner-takes-all networks, sustaining a single bump of activity within a population. The stability of a single bump relies on interactions between an excitatory population with recurrent connections and global inhibition which prevents runaway activity.

Several computational models have been proposed to demonstrate how this circuit can use attractor dynamics to encode heading direction [10, 20–27]. These models all include EPG and PEN cells (or equivalent) as the excitatory component, with reciprocal connections between the two populations forming the bump by propagating activity around the ring network (Fig 1C). For the bump of activity to be maintained there must be recurrent connections back onto the currently active EPG cell. Recent *Drosophila* connectome data has revealed self-recurrent connections between excitatory EPG cells that likely ensure bump persistence [13, 14, 26]. This is in contrast to proposals in previous models where reciprocal connections between EPG cells and PEG cells would maintain the bump [20–23, 25, 27], however PEG to EPG connections are actually very weak [14]. Global inhibition is provided by two further cells types: Δ7 which functionally deliver strong inhibitory input to EPG cells on the ring that are opposite from the location of the bump ([13]; Fig 1D), and GABAergic ring neurons which have characteristic ring-shaped axonal projections enabling inhibition of EPG cells around the EB ([28]; Fig 1E). A combination of both of these inhibitory inputs has been shown to produce a more stable attractor network than either inhibitory source alone [26].

Ring neurons are also responsible for bringing in sensory information such as wind direction [29], polarized light [30], and temperature [31]. All of these sensory cues likely contribute to tethering heading estimation to the environment [32], however the heading representation can also be maintained with angular self-motion information or motor efference copies in the absence of visual information [4]. In this case the head direction estimate is subject to accumulation of error when driven purely by idiothetic cues.

Therefore, understanding how sensory information takes control of the head direction signal is essential for understanding how insects maintain long-term stable estimates of heading. One proposed method of maintaining the head direction estimate is by learning relationships between visual information arriving from visual ring neurons and the EPG cells representing the current heading, such that when a visual scene is revisited the associated heading can be recalled [33, 34].

Ring neuron subtypes ER4d and ER2 have small ipsilateral visual receptive fields that have been characterised in *Drosophila* ([35]; Fig 1B). They have been implicated in a range of visual behaviours [9, 36], and as a population can encode visual features such as size, position and orientation [37]. Visual information reaches ring neurons via the anterior visual pathway [38], from the optic lobe via TuBu cells responsive to many types of visual features in the anterior optic tubercle (AOTu) to ring neurons in the bulb (BU). Similar to cells found in the mammalian visual system, each ring neuron has a receptive field formed of excitatory and inhibitory sub-fields [35].

Previous models of the insect central complex have represented visual input with several methods: simply by mapping the horizontal position of the visual cue (bright vertical bar) onto the 16 EPG cells and stimulating the appropriate PEN cell to move the bump [22, 26]; down sampling the visual scene and applying Gaussian filtering to approximate ring neuron receptive fields, then using the pixel intensity to represent inhibitory ring neuron activity [33]; and representing visual landmarks using multiple ring neuron populations each selectively tuned to the position of a specific landmark in the visual field [25, 27]. Here we present a generalized, minimal spiking neural network model of the insect central complex only including essential connections and cell types, using ring neurons with *Drosophila*-like visual receptive fields to tether the heading estimate to features of a natural visual scene. We use a novel characterisation of the excitatory and inhibitory regions of ring neuron receptive fields that are formed through inhibitory connections between ring neurons (see Methods; [14, 39]).

The aim of this work is to develop a spiking model of the head direction representation in the insect central complex that can be used to explore questions about visual control over the head direction signal. We first ask if simple *Drosophila*-like visual filtering is sufficient to learn a mapping between heading and visual features, and subsequently maintain a heading estimate during rotation and simple translation. When exploring ring neuron visual input to the network, we first challenge the network with simple synthetic stimuli based on previous experimental work to test the model performance compared to known Ring and EPG neuron activity in these analogs of experiments. We then challenge the network with videos of novel natural visual scenes, and investigate how the specifics of natural environments interact with the model to determine performance.

## Materials and methods

The spiking neural network model of the insect central complex was built using PyGeNN, a Python Library for GPU-Enhanced Neural Networks [40] with a timestep of 1 ms. All neuron types were modelled as Leaky Integrate-and-Fire (LIF) neurons with membrane capacitance C*_m_* = 0.2*nF*, resting membrane potential *V*_0_ = *−*70*mV*, reset potential *V_reset_*= *−*70*mV*, spiking threshold *V_spike_*= *−*45*mV*, Membrane time constant *τ_M_* = 20*ms*, and refractory time 2*ms*. Synapses between R2 or R4 neurons and EPG neurons use a custom weight update model (see below). All other synapses are non-plastic so use the standard GeNN ‘StaticPulse’ weight update model. All neurons shape their synaptic input using a single exponential model (‘ExpCurr’ in GeNN) with decay time constants of *τ* = 50 for inhibitory synapses and *τ* = 100 for excitatory synapses.

### Network structure

The connectivity included in this model is based on previous central complex models [23, 25, 26] and supported by recent *Drosophila* connectome analysis [14]. We include only essential cell types and connections in a generalised network that is not species-specific. The model includes five cell types: excitatory EPG and PEN neurons, and inhibitory Δ7, R and Ring neurons (Fig 1B).

The EB is organised into 8 tiles, with each tile subdivided into two wedges containing either a right or left-projecting EPG cell (total = 16; Fig 1A). Left and right-projecting EPG cells target PEN cells in the left and right hemisphere’s protocerebral bridge (PB). The PB contains 16 or 18 glomeruli depending on species, here we include 16 PEN cells, one per glomeruli, each of which is enervated by the equivalent EPG cell (Fig 1A,C). PEN cells return projections back onto EPG cells in a shifted pattern (Fig 1C), targeting two EPG cells one tile around the ring either anticlockwise or clockwise for neurons in the right or left PB respectively, facilitating propagation of activity around the ring. Predictable bump shifts can be induced by stimulating PENa cells [11]. Through their reciprocal connections with EPG cells, PEN cells propagate activity around the ring network and, when left and right PEN cell firing rates are imbalanced, move the bump around the ring. Self-recurrent connections from the the currently active EPG cell ensure that the bump of activity persists. The activity of EPG cells is confined to a single bump by inhibitory inputs from Δ7 neurons [12]. Δ7 cell activity is in turn generated through excitatory inputs from EPG neurons (Fig 1D). The observed connectivity between Δ7 cells and EPG cells is overly complex than would be necessary to contain the bump and likely contributes to other mechanisms, here we use a slightly simplified connectivity pattern suited to a network with 16 EPG cells and 8 Δ7 cells that results in strongest inhibitory input to EPG cells on the opposite side to the bump (Fig 1D,2A). Global inhibition is also provided by one R neuron which is reciprocally connected to all EPG cells (Fig 2A). See Table 1 for specific initial weights.

**Fig 2.**
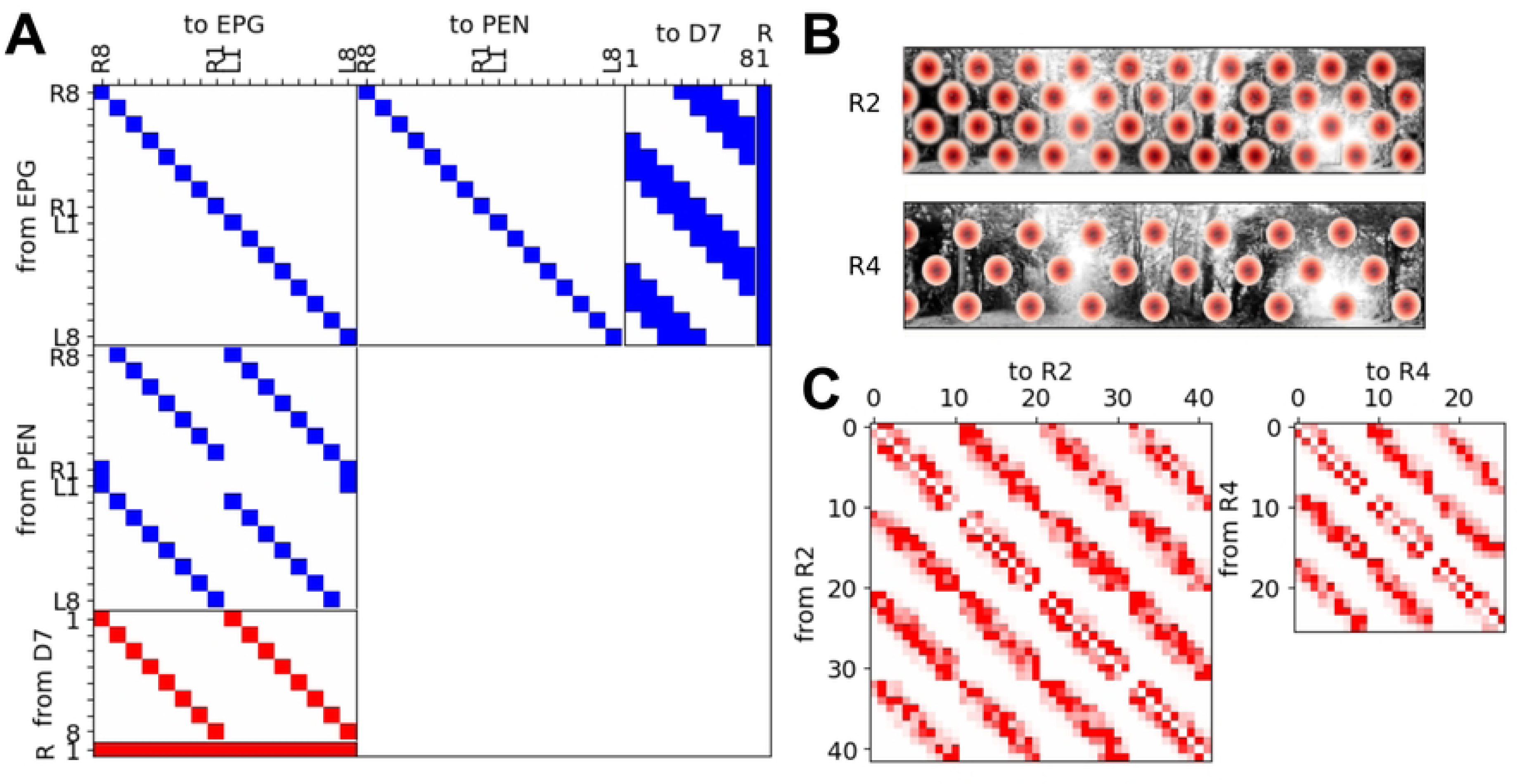
Specific cell to cell connections between populations. (A) Matrix showing connections between EPG, PEN, Δ7 and R cells, following previously determined connectivity (Pisokas et al. 2020, Hulse et al. 2022, Robinson et al. 2022, Chang et al. 2023). Blue and red indicate excitatory and inhibitory connections respectively. EPG *→* EPG (weight 0.02), EPG *→* PEN (weight 0.13), EPG *→* R (weight 0.01), EPG *→* Δ7 (weight 0.05), PEN *→* EPG (weight 0.14), R *→* EPG (weight −1.3), and Δ7 *→* EPG (weight −2.6). (B) R2 (top) and R4 (bottom) ring neuron receptive field centers arranged in hexagonal pattern over the visual scene. (C) Inhibitory connections between ring neurons. These connections decay with distance between field centers with random scaling to produce irregular inhibitory regions of the receptive fields (see Figure). R4 *→* R4 (max weight 0.3) and R2 *→* R2 (max weight 0.3). Not shown R4 *→* EPG (initial random weight between 0 and −0.05), R2 *→* EPG (initial random weight between 0 and −0.05).

**Table 1.** Connection weights between populations.

| Connection | Weight (nA) | Connection | Weight (nA) | Connection | Weight (nA) |
| --- | --- | --- | --- | --- | --- |
| EPG → PEN | 0.13 | PEN → EPG | 0.14 | R2 → EPG | -0.05 |
| EPG → R | 0.01 | R → EPG | -1.3 | R4 → EPG | -0.05 |
| EPG → D7 | 0.05 | D7 → EPG | -2.6 | R2 → R2 | -0.3 |
| EPG → EPG | 0.02 | | | R4 → R4 | -0.3 |

### Getting the bump moving

PEN-a neurons conjunctively encode both head direction and angular velocity [10, 11]. By biasing current input representing angular velocity to one half of the PB we can disrupt the balance between input to EPG cells and drive the bump either clockwise of anticlockwise around the ring [20–23, 25]. Constant angular velocity (corresponding to rotating in one direction at a constant speed) is provided to PEN cells using a GeNN current source model [40] with constant magnitude. Current is supplied with equal positive and negative value to PEN cells in the left and right hemisphere respectively to achieve clockwise rotation and vice versa for anticlockwise rotation. This results in stronger inputs from PEN cells to EPG cells one further clockwise or anticlockwise around the ellipsoid body driving the bump around the ring.

### Ring neuron input to the Ellipsoid Body

The bump position is also manipulated by inhibitory inputs from R2 and R4 ring neurons which bring information about the visual scene. The visual receptive fields of 28 R2 and 14 R4 ring neurons characterised in *Drosophila* [35] are large and distributed unevenly across the visual field (Supplementary Fig 1B), with excitatory and inhibitory sub-fields where the ring neuron activity is modulated up or down when a bright stimulus is presented (Fig 1B). Here we produced synthetic ring neuron receptive fields arranged in a hexagonal grid to evenly tile the 95 *×* 360 degree visual scene (Fig 2B). The hexagonal spacing is set to produce 42 R2 and 26 R4 receptive fields to more closely match the number of each of these cell types identified in the connectome analysis [14]. Each ring neuron has an excitatory RF only, with the inhibitory regions forming through mutual inhibition between ring neurons of the same type [14]; Fig 2C).

The strength of the excitatory receptive field is modelled as a Gaussian function (*σ* = 225) scaled to deliver a current input of between 0 and 0.35 *nA* to the ring neurons. Ring neuron activity is generated by applying the synthetic receptive fields as filters to 95 *×* 360 degree panoramic video frames and using the resulting activation value as current input to ring neurons via a GeNN current source model. The weight of the inhibitory connections between Ring neurons depends on the distance between the RF centers (Gaussian function *σ* = 40), and a random weighting (Fig 2C) which ensures nonuniform inhibitory input and irregular center surround receptive fields as observed in *Drosophila* (Supplementary Fig 1).

This method of generating the inhibitory regions through mutual inhibition is driven by the presence of inhibitory connections between ring neurons highlighted by connectome analysis [14]. The inhibitory connections are mostly specific within ring neuron type [39], and inhibitory regions *Drosophila* ring neuron receptive fields appear similar to the sum across neighbouring excitatory regions of other cells with some random weightings. Using this method to generate synthetic receptive fields allows us to use the full number of visual ring neuron identified by connectome analysis, and to explore how the arrangement of visual inputs to the EB may contribute to targeted learning through selective attention.

### Synaptic Plasticity

Each ring neuron synapses with every EPG cell in the ring (Fig 1F; [28]). To map visual cues represented by the activity of R2 and R4 ring neurons onto headings represented by EPG neurons, Hebbian synaptic plasticity is enabled on these synapses [33, 34]; Fig 1E,F) for the first two rotations of each simulation. As ring neurons are inhibitory, coincident activity at these synapses triggers a change in synaptic weight (*w_i,j_*) towards *w_max_* = 0, weakening the inhibitory synapse. Non-coincident spikes trigger a weight change towards *w_min_*= *−*0.3 increasing the strength of the inhibitory synapse. We implement this using a Spike Timing Dependent Plasticity (STDP) rule based on [41] and implemented as a GeNN custom weight update model. The following weight updates are triggered by ring or EPG neuron spikes:

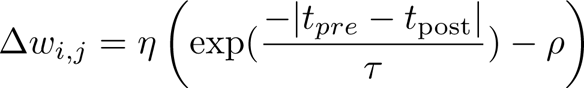

where *t_post_* and *t_pre_* are the time of the postsynaptic and presynaptic spikes respectively, *τ* = 50.0, *ρ* = 0.05, learning rate *η* = 0.008. A parameter search over *η* and *ρ* (Supplementary Fig 3) showed a region of parameter space where combinations of *η* and *ρ* result in low error across all natural scenes (see below). Values were selected from this low error region.

### Visual Stimuli

We first replicated visual stimuli previously used in *Drosophila* experiments [33, 34, 42] and tests of the *Drosophila* receptive fields [9], including a white vertical bar (95 *×* 15 degrees) on a black background, or two white vertical bars (each 95 *×* 15 degrees) with 180 degree separation; and white triangles arranged in a repeating pattern of aligned upright and inverted triangles on a black background or white triangles arranged in a repeating pattern of upright and inverted triangles aligned along their vertical center of mass [42].

We then presented the same network with a total of 33 panoramic videos of natural scenes (Supplementary Fig 2) recorded at various locations across the University of Sussex campus and adjacent Stanmer Park (Supplementary Fig 2). For 23 of these scenes, panoramas were captured as the camera was rotated in place at 10 rotations per minute, and frames were provided as visual input to ring neurons. 17 videos were recorded in densely wooded areas and 6 in open areas in different weather conditions. Videos were captured using a Kodak PIXPRO 360 4k camera and all frames were cropped and resized to 95 *×* 360 px and converted to greyscale. Global histogram equalization was applied for contrast adjustment using the OpenCV library. Rotating datasets were produced by mounting the PIXPRO camera onto a stepper motor programmed to rotate clockwise for 3 revolutions at 10 rotations per minute using an Arduino.

The further 10 scenes were captured using a cable driven camera similar to spidercam or skycam (Supplementary Fig 2) and processed in the same way. The camera was moved through a 200mm radius circle, and the resulting panoramas artificially rotated to maintain heading in line with the direction of movement. The cable-driven camera consisted of a 1 *×* 1 m frame with 4 nema17 47mm stepper motors arranged at each corner, each with a spool of braided fishing line. The fishing line is threaded through eyelets 450mm above the frame attached to transparent acrylic arms then connected at a central point beneath the PIXPRO camera which is counterweighted. By spooling in or out each cable the camera can be moved in 3 dimensions anywhere within the frame. The ground truth position of the camera in a lab environment was recorded using a Vicon motion capture system, that tracks the relative positions of IR reflective markers attached to the camera. Assuming heading is in the direction of movement, the appropriate angle of each frame was estimated from the smoothed ground truth trajectory, and used to artificially rotate panoramic videos after collection. Videos were downsampled to match angular velocity in both the rotation and circling scenes.

## Results

Here we show the behaviour of our spiking neural network model of the insect central complex with inputs from ring neurons with *Drosophila* inspired receptive fields. Using a computational approach allows us to interrogate the network using more complex natural visual input than would be possible during electrophysiological or calcium imaging experiments. To demonstrate the robustness of the network we begin with simple visual stimuli (bright vertical bars) that have been used extensively in experimental work. As we are interested in natural scenes containing conflicting information, we then presented the network with two types of ambiguous stimuli to observe the networks response to conflicting cues (two identical bright vertical bars [33, 34]; series of upright and inverted triangles [42]). Finally we present natural scenes to the network, first rotating the camera on the spot (mimicking how the synthetic visual cues are presented to flies) and then translating through a circle. Most of these experiments follow the same general structure. As we assume this network encodes the mapping between heading and visual features quickly, both the visual input to ring neurons and equivalent angular velocity input to PEN cells were provided for two rotations, and then visual input only for a third rotation (probe trial).

Throughout the results, we demonstrate the ability of the network to robustly maintain a heading estimate in response to a variety of visual stimuli including bright vertical bars and natural panoramic scenes, including stimuli width competing cues. We assess performance based on how closely the center of the bump of EPG activity follows ground truth head angle (see Methods for details). Interestingly, across all experiments the bump of EPG cell activity spans approximately 4-5 cells at any given time, with each of the 16 EPG cells representing approximately 22.5°and a total bump width of between 90°-112.5°– equivalent to 25 - 31.25% of the ring. This is very similar to the spread of activity observed in Calcium imaging of *Drosophila* EPG cells [4].

To measure how closely the model tracked heading using visual features, we compared the estimated heading to the ground truth heading. The estimated heading was found by binning spikes into 16.7 ms bins (time window of one frame) and finding the cells active during each bin and taking the median active EPG cell as the center of the bump, multiplied by the angle represented by each cell (22.5 degrees). The root mean squared error between the estimated heading and ground truth for the final rotation with only visual input available was used to measure model performance over a total of 400 simulations (25 random initial weight spaces between ring neurons and EPG neurons and 16 shifts in initial panorama angle).

### Learning the mapping between simple stimuli and heading

We first replicated three experimental setups used to challenge insects. To observe the learning behaviour of the network when presented with a simple stimulus we used the experimental set-up of [33, 34] and presented the network with a 15 degree wide bright vertical bar on a black background which progressed across the visual scene at the same speed indicated by the angular velocity input to the network (Fig 3A). The bar moved through the visual field for 3 full rotations, ensuring ring neurons with RFs across the full visual scene were coactivated with EPG cells. For the first two rotations, angular velocity information was provided to the network as current injection into PEN cells (Fig 3A, white shaded region), driving the bump around the ring. During the probe trial, the PEN input is turned off and the bump position is driven purely by dis-inhibition from the visual ring neurons (Fig 3A, green shaded region; RMSE 1.17 degrees). Learning at synapses between the ring neurons and EPG cells during the first and second rotation resulted in a weight matrix with clearly visible associations between active ring neurons representing the bar position and heading (Fig 3B).

**Fig 3.**
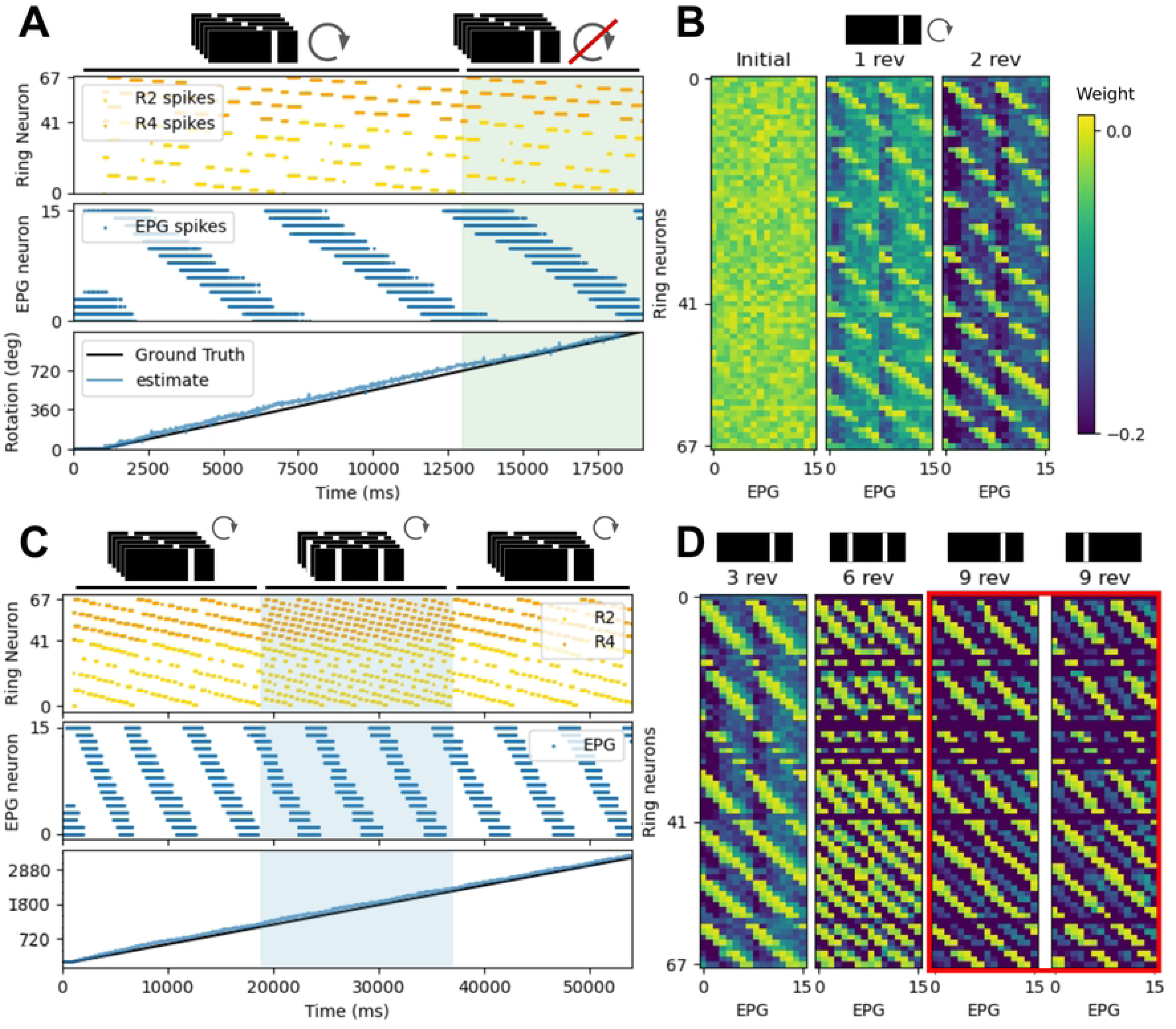
Bump dynamics for vertical bar visual scenes. (A) Raster plots of ring neuron and EPG cell spikes over 3 revolutions with a visual scene of a single bright vertical bar. For two revolutions both visual and angular velocity input are provided to the model. Only visual input provided during the final revolution (green shaded region). The unwrapped ground truth heading and estimated heading (median active EPG cell, see Methods) are also shown. (B) Evolution of the synaptic weight matrix from random initial weights after 1 and 2 revolutions for the one bar visual scene. (C) Raster plots of ring neuron and EPG cell spikes over 9 revolutions. The initial 3 revolutions with a visual scene with a single bright vertical bar. For a further 3 revolutions an ambiguous visual scene with two identical bright vertical bars separated by 180 degrees (blue shaded region). For the final three revolutions the single bar visual scene was returned with the bar at one of the two offsets. A single bump of activity is maintained throughout by ring attractor dynamics. (D) Evolution of the synaptic weight matrix from random initial weights after each visual scene presentation. When two bars are presented, both positions are represented in the weight matrix. The final weight matrix can toggle between two states depending on the offset position of the final vertical bar (red box). Either maintaining the original mapping (3 revolutions vs 9 revolutions left) or remapping to the new bar position (3 revolutions vs 9 revolutions right).

As we are interested in natural scenes with conflicting information we introduced a second conflicting cue (two bright bars separated by 180 degrees; Fig 3C), following the method of [20, 34] to observe changes in the weight matrix. When the network is trained with these ambiguous stimuli, ring attractor dynamics ensure only a single bump of activity is maintained in the ring, although a second representation of the bright bar is learned in the ring neuron to EPG weight matrix (Fig 3D). When returning to a single bar stimulus only one of these two competing representations is maintained (Fig 3D red box), depending on which bar remains. A similar phenomenon has been recorded experimentally in *Drosophila* [33] and these results indicate that the network can successfully form an appropriate, flexible mapping between visual cues represented in the activity of ring neurons and head direction cells, and use this mapping to maintain the bump position in the absence of angular velocity input. Contrary to previous work, our model maintains an accurate representation of the ground truth when presented with ambiguous stimuli rather than skipping over half the population when the cue is revisited [33]. While, in natural scenes with conflicting information, it is important to maintain a consistent representation, this behaviour indicates that visual cues may be less dominant in our model than observed experimentally in *Drosophila*.

Previous work on ring neuron receptive fields has identified patterns of visual stimuli that are discriminable or indiscriminable by ring neurons [42]. We presented the network with example stimuli of these types to observe if the model mimics the experimental observations – even with synthetic ring neuron receptive fields arranged in a hexagonal grid rather than clustered around the horizon to either side of the visual field (Supplementary Fig 1). The two visual cues were made up of a series of upright and inverted triangles, the first of which is known to be discriminable and the other indiscriminable by *Drosophila* receptive fields [9, 42]. The network successfully tracks ground truth head direction during the probe trial when presented with the discriminable stimulus where triangles are aligned at their top and bottom (Fig 4A,B), and is unable to maintain a heading estimate close to ground truth when presented with triangles aligned by their center of mass (Fig 4C,D). Ring neuron activation patterns show the same ring neurons are activated for both the upright and inverted triangles (Fig 4C) resulting in conflicting mappings that in this case prevent the bump movement even when angular velocity information is available. This result demonstrates that the synthetic receptive fields we use have similar discrimination performance as the receptive fields recorded in *Drosophila* [9, 35, 42], and the specifics of ring neuron arrangement and field shape are not essential to this task.

**Fig 4.**
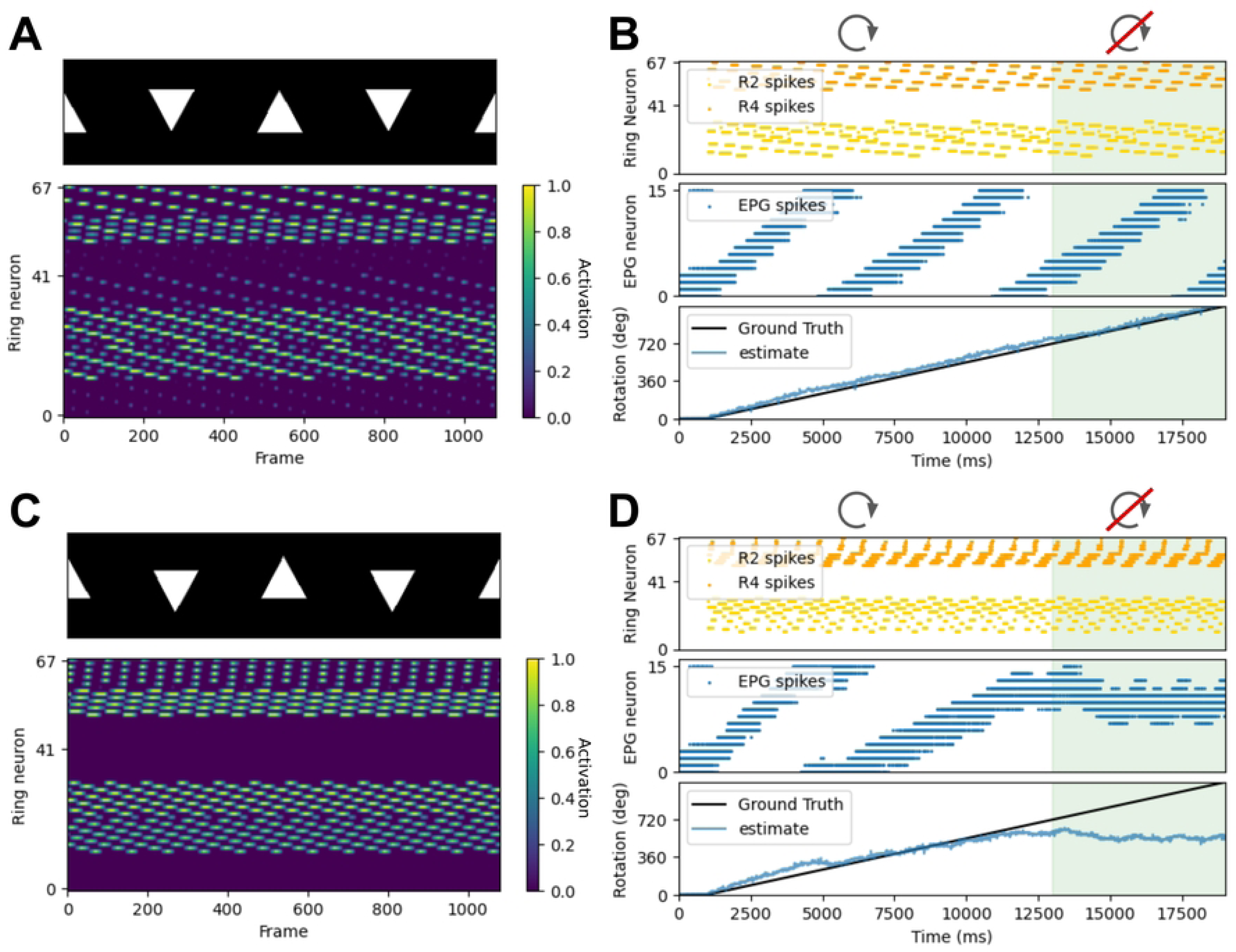
Bump dynamics for vertically aligned and offset triangle stimuli. (A,C) Single frame from each triangle stimulus used by Dewar et al. [9] and associated ring neuron activations across all frames. (A) Triangles arranged in a repeating pattern of aligned upright and inverted triangles, discriminable by Drosophila ring neuron receptive fields [9, 42]. (C) Triangles arranged in a repeating pattern of upright and inverted triangle aligned along their vertical center of mass. Indiscriminable by Drosophila ring neuron receptive fields [9]. (B,D) Raster plots of ring neuron and EPG cell spikes over 3 revolutions, and the unwrapped ground truth heading and estimated heading (median active EPG cell, see Methods). For two revolutions both visual and angular velocity input are provided to the model. Only visual input provided during the final revolution (green shaded area). (D) For the indiscriminable stimulus, the network was unable to form a mapping between ring neurons and EPG cells such that the bump position is maintained once the angular velocity input was removed, as expected if the synthetic receptive fields are a good estimate of the Drosophila ring neuron receptive fields.

### Learning a mapping between natural scenes and heading

Now that we have shown that the network is able to learn a mapping between ring and EPG neurons for simple stimuli, we applied the network to more challenging natural scenes. We presented two types of natural scenes, first mimicking an insect rotating on the spot as in the virtual experiments, then moving in a circular trajectory. Correlation matrices between pixel values or ring neuron activations at each frame of the visual stimuli were calculated by first normalising the frames using OpenCVs normalise function, then finding the Pearson correlation coefficient between pixel values or ring neuron activations for each pair of frames. Over 3 rotations (two with visual and angular velocity input and one probe trial) correlations between frames (Fig 5B,G) and ring neuron activations (Fig 5C,H) were consistent when headings were revisited, showing that the same visual scene results in the same ring neuron activation pattern. However, for some natural scenes, high correlations were also observed between frames and ring neuron activations at additional headings, resulting in multiple peaks, indicating ambiguity or aliasing in the visual scenes (Fig 5G,H).

**Fig 5.**
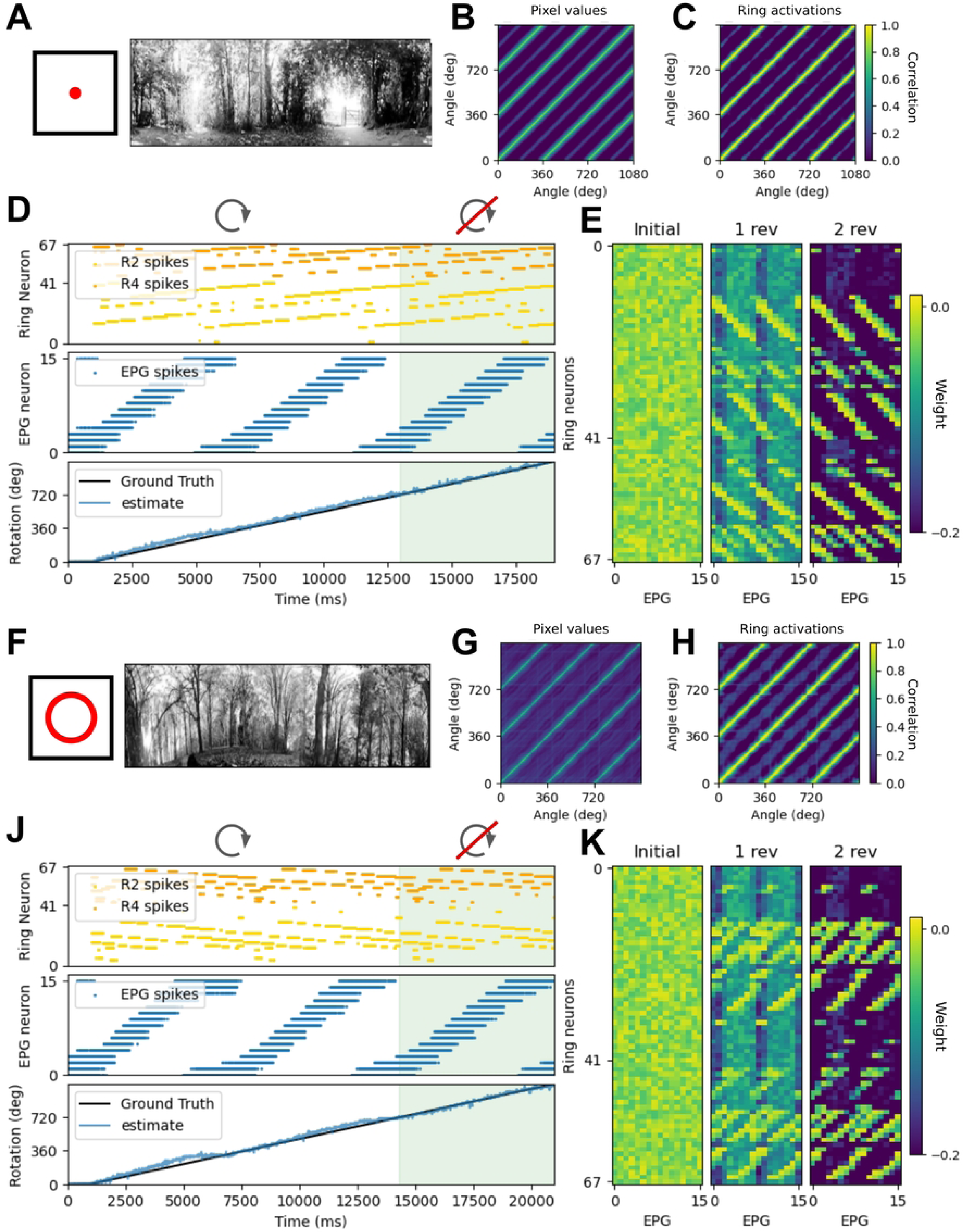
Bump dynamics for revolutions in two natural visual scenes during rotation and translation. (A,F) Single frame from each of the greyscale panoramic videos. Histogram equalisation is applied to each downsampled video (see Methods). (A) Frame video with camera rotation only. F: Frame video with circular camera translation and rotation. (B,G) Correlation matrices showing correlation between each frame for three rotations. Strong correlation is seen between frames at the same rotation angle, but this correlation is not 1 due to natural noise in the video. G: High correlations between ring neuron activations are also seen at intermediate angles indicating ambiguity in the ring neuron representation of the scene. (C,H) Correlation matrices showing correlation between ring neuron activations at each frame for three rotations. Strong correlation is seen between activations for frames at the same rotation angle. (D,J) Raster plots of ring neuron and EPG cell spikes over 3 revolutions, and the unwrapped ground truth heading and estimated heading (median active EPG cell, see Methods). For two revolutions both visual and angular velocity input are provided to the model. Only visual input provided during the final revolution/probe trial (green shaded area). (E,K) Evolution of the synaptic weight matrix from random initial weights after 1 and 2 revolutions.

In both of the examples shown in Fig 5 – one rotating in place and the second moving though a circular trajectory – the network successfully tracked heading during the probe trial. The first example scene was easily discriminable by ring neuron activations, and visual input was sufficient to drive the bump position and maintain an estimated heading close to ground truth (Fig 5D; mean RMSE over 20 simulated seeds and 16 panorama shifts = 0.854 *±* 0.014 degrees; rotation scene 5). For some visual scenes, including the second example, the mapping between ring neurons and heading contained conflicting representations which led to more error in bump movement across repetitions of the simulation (Fig 5J,K; mean RMSE over 20 simulated seeds and 16 panorama shifts = 2.694 *±* 0.214 degrees; circling scene 6). Surprisingly, there was little difference between the two set-ups. We expected that introducing some proximal objects and resulting parallax by placing the camera near the ground or close to trees and undergrowth where there were lots of obstructions would make these scenes especially difficult to track, but heading was tracked robustly.

As the model is capable of tracking heading for both the rotation and circling data sets, we were interested in the robustness of our model and in particular, whether forming a good encoding is more difficult for some scenes. What features of natural scenes might be the most salient or make these scenes easily discriminable? Is a behavioual strategy required to improve performance? It it beneficial to have many ring neurons representing a scene? Do multi-polar ring neurons reduce network performance? To probe these questions we ranked the performance of the model for each natural scene based on the mean RMSE between ground truth and estimated heading for the probe trial over 20 simulated seeds each for 16 starting orientations within the panorama (320 total simulation runs), and divided natural scenes into three classes: those with high variance in the error measure across all simulations (standard error *>* 0.1), those with low variance (standard error *<* 0.1), and scenes with greater than 1% failed simulations (Fig 6A).

**Fig 6.**
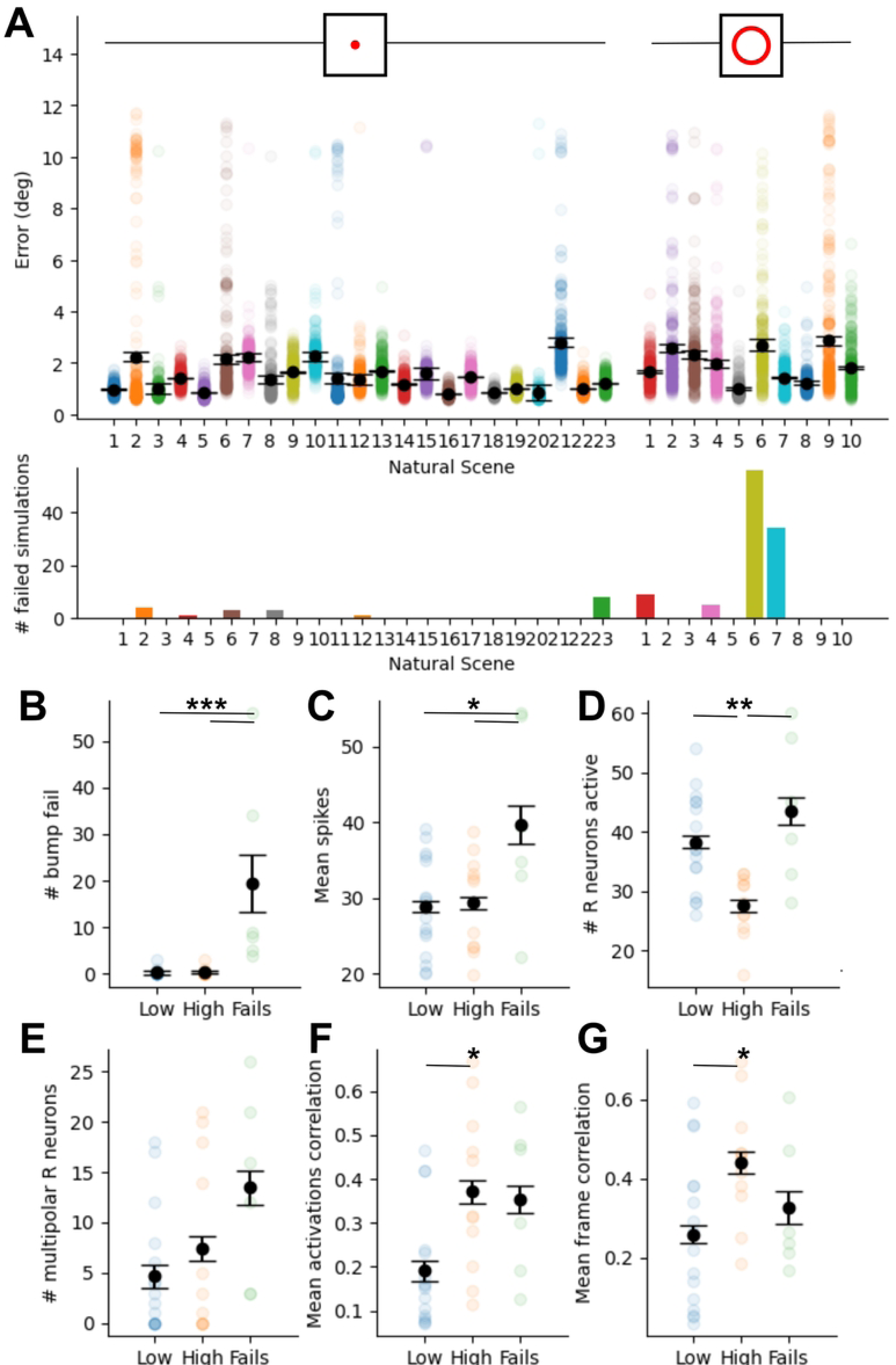
Performance across different natural scenes during rotation and translation. (A) Mean RMSE (degrees) between ground truth and estimated heading during the probe trial with only visual information available, over 20 simulated seeds and 16 shifts of the panorama initial angle *±* bootstrapped standard error (top), and total number of failed simulations for each natural scene, where the bump of activity is lost (bottom). Scenes are separated into camera rotation only (left; 23 scenes), and circular translation and rotation (right; 10 scenes). B-G: Comparisons spiking and image properties between natural scenes with more than 1% failure rate, high or low variance. (A) Number of failures. (C) Mean ring neuron spikes. (D) Total number of ring neurons active. (E) Number of ring neurons with multipolar receptive fields. (F) Mean over ring neuron activation correlation matrix. (G) Mean over video frame correlation matrix. (For all plots *** ANOVA p*<*0.001; ** ANOVA p*<*0.01; * ANOVA p*<*0.05).

We compared these groups across several measures of ring neuron activity and image properties. Scenes in the fail group had on average higher ring neuron mean firing rate than both the low and high variance groups (ANOVA with Tukey’s HSD; Low p = 0.018; High p = 0.037) suggesting an active strategy for adapting peak ring neuron activity may be required to make these scenes more easily discriminable. High variance scenes had fewer total ring neurons active than the low variance and fail group (ANOVA with Tukey’s HSD; Low p = 0.007; Fail p = 0.002) indicating these scenes had features that activated fewer ring neurons, resulting in fewer strong mappings between ring neurons and EPG cells, and overall less power to disinhibit EPG cells and drive bump movement. High variance scenes also showed higher mean values over both the ring neuron activations correlation matrix and the raw pixel value correlation matrix than the low variance group (ANOVA with Tukey’s HSD; mean activations correlation p = 0.019; mean frame correlation p = 0.031), indicating higher correlations of ring neuron activations and pixel values at intermediate headings and directional ambiguity in the visual scene with cells representing multiple parts of the visual scene. In all groups there were examples of natural scenes where individual ring neurons were active for multiple headings resulting in multipolar directional turning. Multipolar tuning is another indicator of directional ambiguity, however the presence of these cells in all groups indicates similar patterns of coactive ring neurons at multiple headings rather than individual ring neurons active at multiple headings that results in more variable performance, and these scenes are less discriminable both at the pixel level and after the large scale filtering by ring neurons.

## Discussion

We have presented a novel spiking neural network model of the insect central complex, incorporating known connectivity [13, 14] between ring neurons [35] and the ellipsoid body compass neurons which allows for the formation of a flexible mapping of visual cues onto headings. The model was able to replicate fly performance and network dynamics across a series of experimental assays with simple visual information such as bright vertical bars [33, 34] or a sequence of triangles [42]. When faced with ambiguous stimuli such as two vertical bars, the network forms multiple representations of the cue and stabilises on one of these when one of the two cues is removed, thus replicating the weight space dynamics proposed by [43]. Similarly, when presented with visual stimuli known to be indiscriminable by *Drosophila* ring neuron receptive fields because of vertically aligned centres of mass [9, 42], the network struggles and can only successfully fix a heading when the shapes are vertically separated, as with flies.

Having demonstrated its plausibility, our goal was to go beyond a proof of concept and challenge the model with complex real-world visual input to see if it could sustain a stable activity bump in the ring attractor. To this end, we used panoramic videos that include natural environmental variations (due to wind and lighting changes) and camera movement. We found that the information encoded by the simple visual filtering of a small number of ring neurons with *Drosophila*-like receptive fields is sufficient to robustly maintain an estimate of heading despite the complex visual input. In the following sections we discuss the implications of these results.

### Capturing the ring neuron to central complex circuit

As our focus was on how visual ring neuron inputs influence the network rather than specifically how the bump is formed or maintained, we constrained the model to the essential underlying architecture and connections required to produce the expected ring attractor dynamics, whilst using cell types and connections from up-to-date connectome data [13, 14]. Notably, PEG cells were excluded which were instrumental in previous work for maintaining bump persistence [20–23, 25, 27], because their function was replaced by direct self-recurrent connections between EPG cells identified in the connectome analysis [14]. As the bump properties remain similar in our model, we now need to look to experiments as to whether PEG cells and self-recurrent EPG activity are redundant mechanisms or whether each has it’s own influence over the circuit.

Secondly, to explore the transformation from visual input to neural code via the ring neurons, we interrogated the connectome to see if there were connections that could create the receptive fields previously described by [35]. We model each ring neuron with only an excitatory receptive field and propose that the function of the previous unexplained mutual inhibitory connections between the ring neurons [14, 39] is to form the functional inhibitory regions recorded in *Drosophila* [35]. Mutual inhibition prevents neurons with overlapping receptive fields from representing the same cue, helps selecting more prominent visual cues and suppresses noise arising from less activated ring neurons. Although here we only include inhibition within ring neuron type, some connectivity between ring neuron types has also been identified, which could enforce a suppression hierarchy when more useful sensory information (such as polarized light or sun position as represented by other classes of ring neuron) win out over less strong cues [14]. These ideas provide interesting testable predictions for future work.

In simplifying the circuit we made several assumptions that limit the biological completeness of the model, but importantly the simplifications are not crucial to the questions explored. For instance, cell types with no clear function and ring neuron subtypes carrying information about other sensory modalities were excluded from the model. We also used a single neuron to provide global inhibition to the network, whilst in reality, this is likely to be provided by a population of ring neurons which may be the same cells as those bringing sensory information into the network [26]. This dual role of ring neurons aligns with the experimental data where the activity of ring neurons is modulated up or down with respect to stimuli passing through the excitatory and inhibitory regions of the receptive field [35]. This raises questions about how this type of activity would impact the mapping between visual cues and the heading estimate. To incorporate this type of ring neuron coding, an alternative learning rule may be required, as we used a simple learning rule that does not take into account the order of spikes or their precise timing.

We made further assumptions regarding the arrangement of ring neuron visual receptive fields. Hexagonally arranged ring neuron receptive fields allow the network to use all available cues across the visual field, and are appropriate for keeping track of large, low-frequency visual cues [36]. Robust model performance shows the arrangement of ER2 and ER4d ring neuron receptive fields as they are observed in *Drosophila* (clustered along the horizon to the left and right of the visual field; [35]) is not essential for tracking heading in this particular task. However, this arrangement may be useful for tracking natural objects or specific environmental features. It may also be the optimal position for extracting visual information relevant to the control of avoidance or attraction behaviours [44]. Other ring neuron subtypes sensitive to polarized light have receptive fields in the upper third of the visual field [45]. Recent work also calls into question the shape of ring neuron receptive fields reported by Seelig and Jayaraman [35]. Based on the dendritic arrangement of upstream MeTu cells in the medulla of the optic lobe [45], the receptive fields of ER4d neurons may be long vertical stripes across the visual field rather than the classic centre-surround type. However centre-surround receptive fields could be achieved for different upstream morphology with a more complex inhibitory connectivity between ring neurons.

Indeed, inhibition is important in the model. The inhibitory connections between ring neurons leads the model to track visual heading with a simple form of selective attention. Varying the strength of mutual inhibition between ring neurons, or additional visual processing upstream in the anterior visual pathway, could also provide more structured visual input to the network to make salient features more prominent or task specific.

### Towards an understanding of the circuit in real-world insect behaviour

In the real world, insects need to know when to learn, but also when not to learn. For our model, the average error across all scenes remained small (below 2 degrees) regardless of whether learning remains on or off during the probe trial, although performance improved when learning was turned off (Supplementary Fig 3B). However, in insects learning rates are not static. For instance, learning at the ER*→*EPG synapses is modulated by dopamine signalling which actively increases synaptic plasticity when angular velocity is large. This prevents over-learning of head directions at slow speeds that otherwise would be over-sampled [46] and would be a valuable future addition to the model.

It is also interesting to question what features of the visual scene insects attend to. With its simple implementation of biomimetic visual processing focused on luminance-defined cues, the network currently tracks areas of the scene with the highest intensity of light such as the sun position or visible areas of sky between trees. Ring neurons have been previously shown to respond to bright sun-like objects or bright areas of a sky gradient [38] which are sufficient for stabilizing the head direction signal in flies [47]. However, it is important to note that although our model can use the sun to track heading on a sunny day, it is not required, and the network was also able to track heading for the same natural scenes on overcast days (Supplementary Fig 2).

Furthermore, the noise inherent in the videos of natural scenes was almost entirely removed after filtering the input by ring neuron receptive fields. However, while attention may be driven by one overriding cue - light level in this instance - heading tracking still requires a population of ring neurons. By comparing natural scenes that are easy or difficult for the network to track, it is clear that rather than headings being represented by the single most active ring neuron, headings are encoded by a combination of ring neurons, each of which may be included in other combinations for other headings. This is evidenced by the appearance of multipolar cells in all performance groups where individual ring neurons are active for multiple headings.

Natural scenes contain more information than just light intensity. Motion-defined cues are also represented by ring neurons, specifically ER4d cells which are sensitive to both motion and brightness [48]. Extending the model to include more of the visual pathway would allow us to build more complex representations of visual scenes and observe how this impacts the network performance. However, it is important to remember that visual information is not the only sensory information available to the fly to keep track of heading, and there are further ring neuron types, which bring in information about other sensory modalities to the central complex [29–31, 38]. This raises the question, how these different sensory modalities are integrated to produce the most accurate estimate of heading. A total of 46 EPG cells were identified in the *Drosophila* connectome [14], of which we currently include 16. This is almost enough cells to have 3 complete rings. This raises the intriguing possibility of having a multi-ring model in which multiple ring attractors act as redundancy in the network or alternatively, each ring may receive different sensory information, brought in by subtypes of ring neurons, and integrating over these inputs would produce a multi-sensory heading estimation.

### Implications for robotic applications and future experiments

Having shown that the model can function robustly with naturalistic input, this raises interesting implications for autonomous robotics systems that would need to maintain estimates of pose or heading in complex natural environments [49, 50]. Studying biological systems in this way highlights the potential for low-energy dedicated biomimetic circuitry, which could be plausibly implemented on neuromorphic circuits for low-power autonomous robotics applications [51–53]. Insect-inspired circuits are a particularly promising avenue for this approach as, while they solve the same problems as mammals (like tracking heading with compass neurons), they do so with far fewer neurons. Robots can also be valuable test-beds for biological hypotheses [54] and we have previously used them to elucidate models of mammalian navigation [55].

One of the most interesting implications of our model is its potential to scale. For example, one could investigate duplicating the ring of EPG neurons for multi-sensory integration, with each ring dedicated to a different behavioural task or sensory modality. Furthermore, different insects are specialists for different behaviours, and although we know the broad architecture of the central complex is conserved across species [56] there must be differences in the circuit that are dedicated to specialist behaviours. Because the majority of experimental data – particularly optogenetics and connectome studies – have focused on *Drosophila* [11, 12, 14, 30, 35, 38, 43, 45], the model presented in this work and most other previous models are biased towards the *Drosophila* circuit [10, 22, 25, 26]. However the real power of computational approaches is that they allow us to explore questions about other insect species and investigate how behavioural specialisms may come about from this conserved circuitry.

## Supporting information

**S1 Fig. Ring neuron receptive fields.** A: 42 synthetic R2 receptive fields 95 x 360 pixels. Excitatory regions (red) of all of all other R2 cells multiplied by a random connection weight are subtracted from each R2 cell to produce the inhibitory region (blue). B: 14 R2 and 7 R4 averaged Drosophila receptive fields (Dewar et al 2017). 112 × 252 pixels for R2 neurons and 88 × 198 pixels for R4d. C: 26 synthetic R4 receptive fields 95 x 360 pixels.

**S2 Fig. Natural scenes.** One example frame from 33 panoramic videos of natural scenes. Videos were captured at various locations across the University of Sussex and adjacent Stanmer Park. (top) 23 rotation only videos. For 3 locations examples were captured in open areas on both sunny and overcast days (total 6 panoramas). The remaining examples include trees either in a woodland or campus setting which occlude some or all of the sky. (bottom) 10 circling videos. A photograph of the spidercam showing 4 cables connected to the camera assembly. Cable lengths are changed by winding or unwinding the cables from a spool using stepper motors, in order to move the camera in 3 dimensions.

**S3 Fig. Varying the Learning parameters.** (Left) Logged mean error for the final rotation for all simulations across rotation only natural scenes, for combinations of parameters learning rate (*η*) and *ρ* (see Methods for learning rule). Parameters *η* = 0.01 and *ρ* = 0.06 were selected from the dark blue region of low error (white star). Colourbar indicates logged mean error over all natural scenes in degrees. (Right) Average error across all natural scenes when learning remains on or turned off during the probe trial (T-test p = 0.001).

## Acknowledgments

None

